# Setting off the Alarms: *Candida albicans* Elicits Pro-Inflammatory Differential Gene Expression in Intestinal Peyer’s Patches

**DOI:** 10.1101/511667

**Authors:** Navjot Singh, Heather C. Kim, Renjie Song, Jaskiran K. Dhinsa, Steven R. Torres, Magdia De Jesus

## Abstract

*Candida albicans* has been associated with a number of human diseases that pertain to the gastrointestinal (GI) tract. However, the details of how gut-associated lymphoid tissues (GALT) such as Peyer’s patches (PPs) in the small intestine play a role in immune surveillance and microbial differentiation, and what mechanisms PP use to protect the mucosal barrier in response to fungal organisms such as *C. albicans*, are still unclear. We particularly focus on PPs as they are the immune sensors and inductive sites of the gut that influence inflammation and tolerance. We have previously demonstrated that CD11c^+^ phagocytes located in the sub-epithelial dome (SED) within PPs sample *C. albicans*. To gain insight on how specific cells within PPs sense and respond to the sampling of fungi, we gavaged mice with *C. albicans* strains ATCC 18804 and SC5314 as well as *Saccharomyces cerevisiae*. We measured the differential gene expression of sorted CD45^+^ B220^+^ B-cells, CD3^*+*^ T-cells, and CD11c^+^ DCs within the first 24 hrs post-gavage using nanostring nCounter® technology. The results reveal that at 24 hrs, PP phagocytes were the cell type that displayed differential gene expression. These phagocytes were both able to sample *C. albicans* and able to discriminate between strains. In particular, strain ATCC 18804 upregulated fungal specific pro-inflammatory genes in CD11c^+^ phagocytes pertaining to innate and adaptive immune responses. Interestingly, PP CD11c^+^ phagocytes differentially expressed genes in response to *C. albicans* that were important in the protection of the mucosal barrier. These results highlight that the mucosal barrier not only responds to *C. albicans*, but also aids in the protection of the host.

**Importance:** The specific gene expression changes within PPs that send the warning signals when encountering fungi, and how PPs can discriminate between innocuous *S. cerevisiae* or different strains of *C. albicans* during early stages of sampling, have not been elucidated. Here we show that within the first 24 hours of sampling, CD11c^+^ phagocytes were not only important in sampling, but they were the cell type that exhibited clear differential gene expression. These differentially expressed genes play important dual roles in inflammation, chemotaxis, and fungal specific recognition, as well as maintaining homeostasis and protection of the mucosal barrier. Using nanostring technology, we were also able to demonstrate that PPs can distinguish between different strains of *C. albicans* and can “set off the alarms” when necessary.

## Introduction

Although a commensal organism and part of the natural human mycobiome, colonization of *Candida albicans* in the gastrointestinal (GI) tract has been associated with a number of diseases such as gastric ulcers, familial Crohn’s disease and Hirschsprung-associated enterocolitis (1-5). It has been suggested that *Candida* colonization yields a continuous low-level inflammatory state in the GI tract that ultimately delays healing and contributes to further colonization (1, 6). Several studies in murine models have demonstrated that *C. albicans* alone is unable to evoke sufficient levels of inflammation to allow for colonization in the GI tract (1, 7-10). However, as a commensal and opportunistic organism, *C. albicans* has demonstrated the ability to exploit an inflammatory state in mouse models using dextran sodium sulfate (DSS) to induce inflammation and antibiotics to deplete the microbiota (1, 7-10).

There is still more work to be done to elucidate details of *C. albicans*’ relationship with the microbiome, as well as the mechanisms through which it takes advantage of the state of the GI tract. In particular, more information is needed on how the host’s gut-associated lymphoid tissues (GALT) initially sense *C. albicans*, and how these tissues ultimately coordinate the specific immune response based on what they have sampled. Specifically, it is unknown how GALT such as ileal Peyer’s patches (PPs), which are considered the inductive sites of intestinal immunity, detect *C. albicans*. Research is needed to explore the possibility that specific gene expression changes within PPs elicit warning signals when encountering fungi. Also, it remains unclear whether PPs discriminate between innocuous *S. cerevisiae* or different strains of *C. albicans* during early stages of sampling.

As the inductive sites of intestinal immunity, PPs are the primary sites through which particulate antigens and microbes are sampled (11). The microarchitecture of PPs is quite unique, as they are interspersed along the antimesenteric side of the ileum and can be seen with the naked eye as aggregate lymphoid structures (11, 12). Each PP contains multiple follicles that are individually equipped with an epithelial barrier called the follicle-associated epithelium (FAE). The FAE contains specialized enterocytes called microfold cells (M-cells) that allow the controlled transcytosis of luminal antigens and microbes. Below the FAE in the sub-epithelial dome (SED), there is an extensive network of at least five subsets of mononuclear phagocytes and a B-cell-rich germinal center (GC) that also contains follicular dendritic cells flanked by T-cell-rich interfollicular regions. In concert, these cells all play an important role in sampling and immune surveillance (11, 13). Because PPs can be a very delicate and technically difficult tissue to work with, many gut models have used cell lines such as the Caco-2 and FaDu to understand sampling and transcytosis (14, 15). Recent work by Rast et al. used the human FaDu epithelial cell line as a model of mucosal epithelial barriers to test early interactions with *C. albicans* (15). Using transcriptional profiling, they found that these cells could discriminate between commensal and pathogenic interactions of *C. albicans* in a dose-dependent manner (15). In our previous studieswe addressed *in vivo* whether PPs could sample *C. albicans* and *C. tropicalis,* and demonstrated that both are sampled within the first twenty-four hours post-gavage(16). We found that *C. albicans* strains ATCC18804 and SC5314, as well as *C. tropicalis*, were able to cross the intestinal epithelial barrier, and that uptake was partly dependent on M-cells. These findings were supported by Albac et al., who demonstrated *C. albicans* strain SC5314 uses M-cells as a portal of entry into the intestinal barrier (16, 17). *C. albicans* was able to preferentially invade M-cells via actin-mediated endocytosis rather than active penetration (16, 17). In addition to its ability to cross the epithelial barrier, we also found that *C. albicans* was sampled by a subset of SED CD11c^+^ phagocytes that expressed the C-type lectin Langerin (16).

Earlier studies by Bonifazi et al. showed that PP dendritic cells (DCs) had functional plasticity and played a role in both inflammation and tolerance in the presence of *C. albicans in vivo*. Interestingly, the immune response that was elicited depended on whether *C. albicans* was in a yeast or hyphal form (16, 18). *C. albicans* in the yeast form promoted the activation of *Rorc*^+^17-producing CD4^+^T cells and *Gata3*^+^IL-4/IL-10-producing CD4^+^T cells with minimal activation of *tbet*^+^IFN-γ-producing cells, an immune response that resulted in a limited control of fungal growth (16, 18). In contrast, hyphae-pulsed PP-DCs promoted the activation of Th1/Treg cells, suggesting the activation of a protective tolerogenic response to *C. albicans* that accounted for the control of fungal growth and long-term survival (16, 18). They concluded that *C. albicans* is a commensal that is capable of balancing inflammation and tolerance at mucosal sites through DCs.

Following up on the studies done by Bonafazi et al., we were not only able to specifically isolate PP CD11c^+^ phagocytes, but also B220^+^ B-cells and CD3^+^ T-cells post-gavage with *C. albicans* (12, 19, 20). mRNA from these cell subsets was isolated for transcriptional profiling as previously described (12). Nanostring nCounter technology uses a novel digital color-coded molecular barcode technology to measure gene expression based on the counts of the target RNA (12). We used the nanostring nCounter^®^ mouse Pan Cancer Immune profiling panel that profiles over 770 murine immune system-related genes. Unlike RNA sequencing (RNAseq), the targeted transcriptomics using nanostring technology do not require library preparation and complex data processing. Nanostring technology is also a quite flexible, as it has also been used as a gene expression profiling platform to probe *C. albicans* during infection (21). To further understand the signals within PPs that sense and respond to *C. albicans* within the first twenty-four hours of sampling, we used nanostring technology on the isolated PP CD11c^+^ phagocytes, B220^+^ B-cells, and CD3^+^ T-cells to determine changes in gene expression.

To identify whether PPs could differentiate between *C. albicans* strains, we chose the commonly used SC5314 (Clade 1) and ATCC 18804 (Clade 5), with *Saccharomyces cerevisiae* as a control. Previous studies by Marakalala and colleagues showed that innate immune receptors such as the C-type lectin receptor Dectin-1 were essential for controlling infection *in C. albicans* strain SC5314, though not required for *C. albicans* strain ATCC 18804 (22). They also found that the differences in Dectin-1 mediated recognition *in vivo* were linked to cell wall composition and architecture. Additionally, strain ATCC 18804 was only recognized by Dectin-1 when mice were treated with the antifungal caspofungin that changed the cell wall architecture (22). The marked differences in innate immune responses between these two strains made them the ideal candidates to use as probes for PPs.

Using transcriptional profiling of 770 genes by nanostring, here we show that PP are able to sense fungi within the first twenty-four hours post-gavage with the yeast form of *C. albicans* strains ATCC 18804 and SC5314, as well as *Saccharomyces cerevisiae*. Specifically, CD11c^+^ phagocytes were found to be the cell type that exhibited differential gene expression within this time frame. We found that *C. albicans* stimulated genes involved in inflammation, chemotaxis and fungal specific recognition. As in the studies by Bonifazi et al., we observed activation of genes that played a role in maintaining homeostasis and protection of the mucosal barrier, such as Interleukins (IL) IL-22 and IL-23. However, *C. albicans* did not have to be in its hyphal form to accomplish activation of these genes involved in the repair of mucosal barriers. When comparing *C. albicans* strains SC5314 and ATCC 18804, we found marked differences in the gene expression profiles of CD11c^+^ phagocytes. Strain SC5314 stimulated genes involved in innate immunity, such as lectins Chitinase-like protein-3 (Chil 3) and Dendritic Cell-Specific Intercellular adhesion molecule-3-Grabbing Non-integrin (DC-SIGN), as well as complement. Strain ATCC 18804 elicited pro-inflammatory fungal-specific cytokines, including IL-17a, and other chemokines and cytokines involved in the recruitment of monocytes and neutrophils, as well as prostaglandin-endoperoxide synthase-2 (Ptgs2), a key enzyme in the biosynthesis of prostaglandin-2, various complement proteins involved in the membrane attack complex (MAC), and S100 Calcium Binding Protein A8 (S100a8), a protein that is also known as as an alarmin. To our knowledge, ours is the first study to use large-scale transcriptional profiling of PPs in response to *C. albicans*. We found that this approach to probe PPs cell subsets was quite informative and can be applied universally to study not only how PPs detect fungi in the intestinal mucosa at different time points, but also how they recognize a variety of organisms.

## Results

### PP SED CD11c^+^ phagocytes sample *C. albicans* strains SC5314, ATCC 18804 and *S. cerevisiae*, and can be isolated to assess gene expression profiles

Because of the documented differences in infection models with *C. albicans* strain SC5314 and ATCC 18804, we tested whether these strains, as well as the *S. cerevisiae* control, were sampled equally by PPs CD11c^+^ phagocytes (22). Mice were gavaged with 1×10^8^ organisms and PPs were harvested 24 hrs post-gavage. The results reveal that all strains, including *S. cerevisiae*, were comparably sampled by CD11c^+^ phagocytes located in the SED of the patch **(Fig. 1).** Since sampling was detected in PPs, we made single-cell suspensions from pooled PPs 24 hrs post-gavage. The cells were sorted to specifically isolate CD45R/B220^+^(CD3^-^/CD11c^-^) B-cells, CD3^+^(CD45R/B220^-^/CD11c) T-cells, and CD11c^+^(CD45R/B220^-^/CD3^-^) phagocytes as previously described (12, 19, 23). During the sorting of PP cells from each of the treatment groups, we carefully selected the gates denoted in green **(Fig. S1A-E for all groups)**, ensuring that there was no cross-contamination from each of cell types of interest.

**FIG 1.**
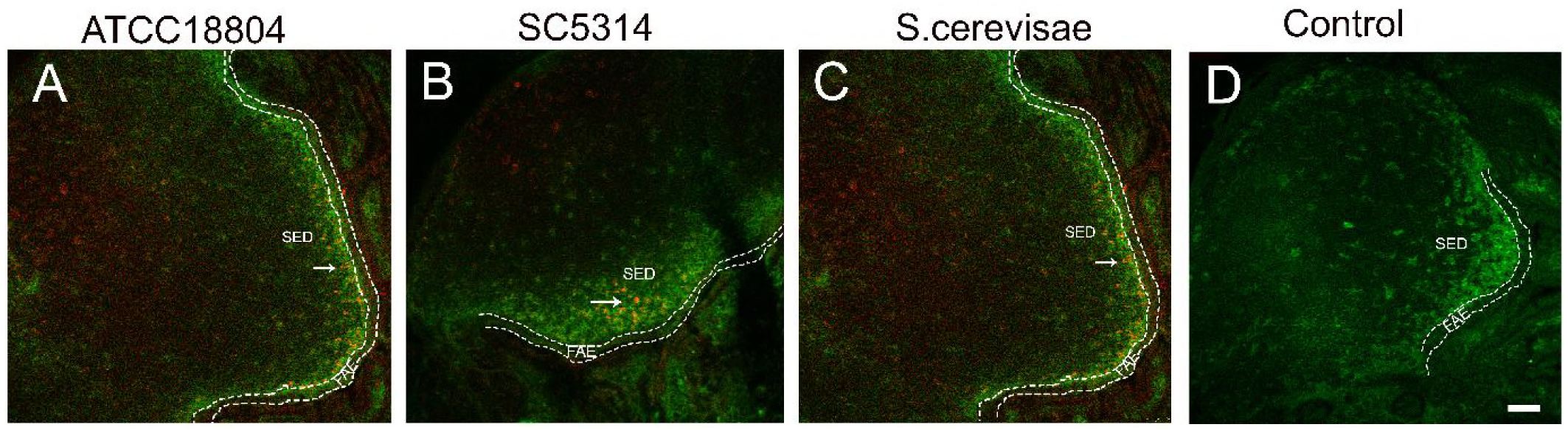
PP SED CD11c^+^ phagocytes sample *C. albicans* and *S. cerevisiae* 24 hours post-gavage. Fluorescently labeled *C. albicans* strains ATCC 18804, SC5314 and *S. cerevisiae* (red) are gavaged and PP are harvested 24 hours later for cryosectioning. PPs were cyosectioned, immunostained with an anti-CD11c^+^ antibody (green) and viewed by confocal laser scanning microscopy. The follicle-associated epithelium (FAE) is highlighted by the dashed lines. Arrows indicate CD11c^+^ phagocytes in the sub-epithelial dome (SED) that have sampled the indicated strain. Scale bar is 50 µm.

### Differential gene expression in CD11c^+^ DCs shows that PPs can recognize fungi and can distinguish between them

To determine if the isolated PP immune cells could detect the presence of *C. albicans*, and whether they could distinguish between *C. albicans* strains ATCC 18804 and SC5314 or *S. cerevisiae*, we assessed the transcriptomic profiles of each cell type using the nanostring nCounter® mouse Pan Cancer Immune profiling panel, which profiles over 770 murine immune system-related genes. The advantage of nanostring technology is that it allows for the detection of differential gene expression with manageable data sets. Statistically significant differential gene expression was determined by analyzing RNA raw counts with DEseq2. For a transcript to be considered differentially expressed in our data set, it had to meet the stringent requirements of having at least 2-fold up or downregulation and a *P*-adjusted value of <=0.05 (12). Principle components analysis of the data sets revealed that each cell type clustered distinctly based on the gene expression profiles, and the segregation was cell type-specific. However, significant changes in gene expression were only observed only in CD11c^+^ phagocytes, confirming that these cells can actively sample fungi and are the next line of defense after crossing the epithelial barrier, as we had previously documented (13) **(Fig. 2)**.

**FIG 2.**
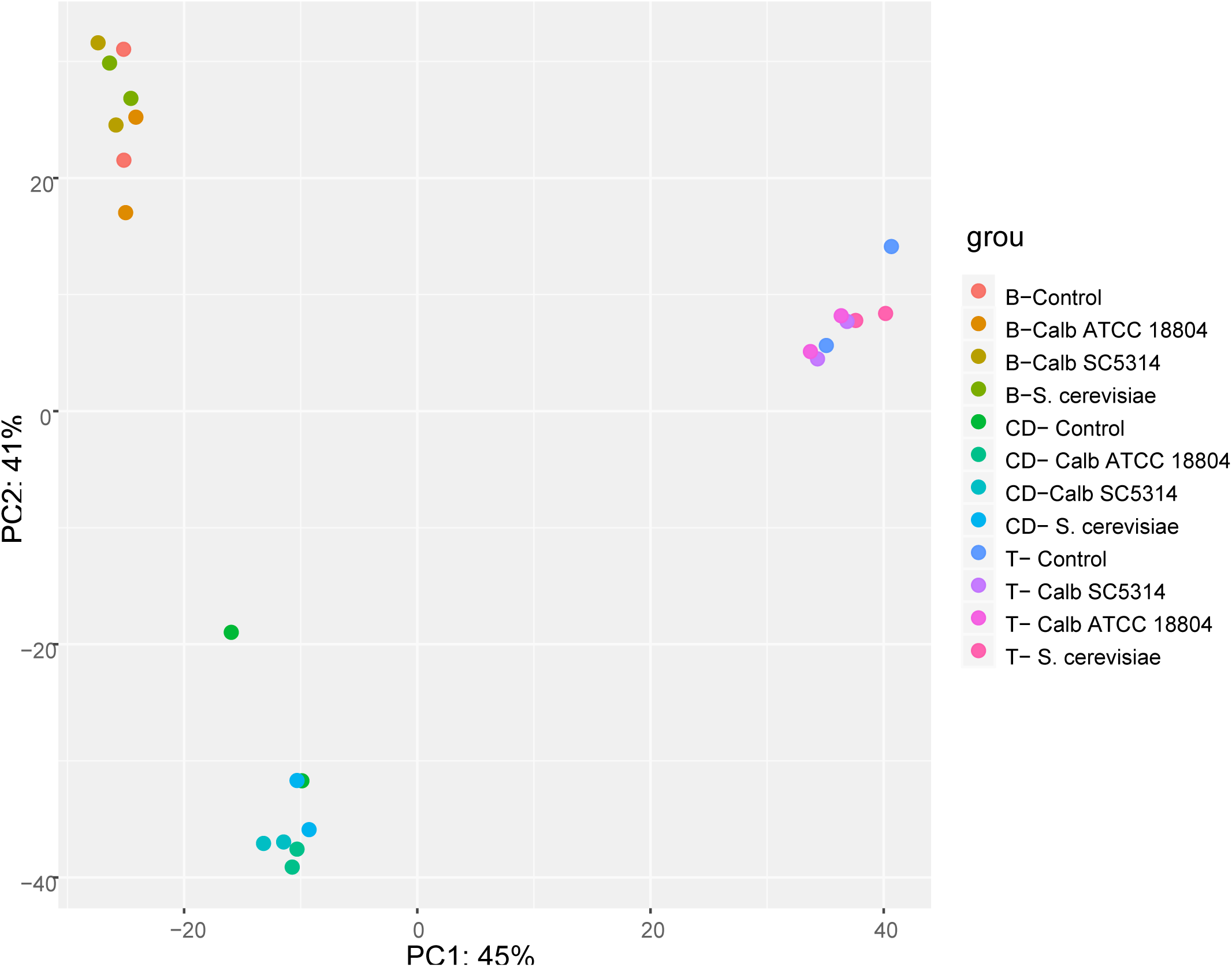
Principal Component Analysis (PCA) of the gene expression levels from PPs that sampled *C. albicans* ATCC18804, *C. albicans* SC5314, *S. cerevisiae,* and untreated control 24 hours post gavage. Nanostring gene expression profiling was done using FACS-sorted B-lymphocyes, T-lymphocytes, and CD phagocytes and analyzed using DESeq2. PCA showed that different cell types are main source of variability in gene expression.

**Table 1** shows the fold values for all genes that met both criteria for determining differential expression. We found that SC5314, ATCC 18804, and *S. cerevisiae* all activated the expression of IL-22 and IL-23 genes in CD11c^+^ phagocytes **(Fig. 3A)**. These cytokines are known to play important roles in maintaining barrier integrity and in antifungal immunity (24). In particular, IL-22, is a key cytokine that has been demonstrated to increase gut epithelial and immune barrier functions by inducing antimicrobial activity, as well as stimulating tissue-damage protection and tissue repair and remodeling, suggesting that PPs recognize all three microbes as a potential threat (25). Fold changes for IL-22 were 18.4-fold upregulation for strain SC5314, 28.1-fold upregulation for *S. cerevisiae*, and 91-fold upregulation for strain ATCC18804. IL-23r is an important pro-inflammatory cytokine receptor that plays a role in antifungal immunity that can drive Th17 responses (25, 26). Fold changes for IL-23r were found to be 2.5-fold upregulation for strain SC5314, 4.2-fold upregulation for *S. cerevisiae*, and 4.9-fold upregulation for strain ATCC18804 **(Fig. 3A)**.

**Table 1:** Differential Gene Expression of PP CD11c^+^ phagocytes

| *C. albicans* ATCC 18804<br>(Clade 5) |  |  | *C. albicans* SC5314 (Clade 1) |  |  | *S. cerevisiae* M5958 |  |  |
| --- | --- | --- | --- | --- | --- | --- | --- | --- |
|  | *Fold*<br>*Change* | *Padj* |  | *Fold*<br>*Change* | *Padj* |  | *Fold*<br>*Change* | *Padj* |
| SHARED DIFFERENTIALLY EXPRESSED GENES |  |  |  |  |  |  |  |  |
| **Il22** | 91.4605 | 5.94E-19 | **Il22** | 18.4034 | 2.68E-07 | **Il22** | 28.1269 | 1.23E-09 |
| **Il23r** | 4.9425 | 2.36E-05 | **Il23r** | 2.5411 | 5.78E-02 | **Il23r** | 4.2218 | 6.39E-04 |
| **Fn1** | 16.6482 | 5.61E-05 | **Fn1** | 41.112 | 1.19E-08 |  |  |  |
| **Ifitm1** | 6.9826 | 5.61E-05 | **Ifitm1** | 5.0496 | 2.92E-03 |  |  |  |
| **Il1r1** | 5.1008 | 1.05E-04 |  |  |  | **Il1r1** | 4.3844 | 2.82E-03 |
| **Il1b** | 5.3101 | 1.17E-04 | **Il1b** | 3.525 | 1.35E-02 |  |  |  |
| **Ifitm2** | 2.6592 | 1.74E-02 | **Ifitm2** | 2.9111 | 1.03E-02 |  |  |  |
| **Cxcl3** | 14.9641 | 2.92E-02 | **Cxcl3** | 29.0585 | 4.74E-03 |  |  |  |
| **Tnfsf11** | 3.8776 | 2.90E-02 |  |  |  | **Tnfsf11** | 5.8876 | 4.57E-03 |
| **Rorc** | 3.0706 | 4.27E-02 |  |  |  | **Rorc** | 3.6993 | 5.25E-02 |
| **Csf3r** | 3.7311 | 2.33E-02 | **Csf3r** | 4.4431 | 1.03E-02 |  |  |  |
| **Csf1** | 4.7156 | 1.35E-02 | **Csf1** | 5.4264 | 7.65E-03 |  |  |  |
| DIFFERENTIALLY EXPRESSED GENES IN INDIVIDUAL STRAINS |  |  |  |  |  |  |  |  |
| **Vegfa** | 12.7209 | 7.71E-05 | **Chil3** | 31.5328 | 7.67E-05 |  |  |  |
| **Ctla4** | 6.4072 | 7.71E-05 | **Ccl24** | 25.0114 | 1.35E-02 |  |  |  |
| **Lif** | 6.4483 | 3.02E-04 | **CD209e**<br>**(DC-**<br>**SIGN)** | 17.5189 | 1.46E-02 |  |  |  |
| **Cd14** | 6.7663 | 4.23E-03 | **Cxcr2** | 5.5573 | 2.02E-02 |  |  |  |
| **C4b** | 4.302 | 6.31E-03 |  |  |  |  |  |  |
| **Thbs1** | 10.9624 | 6.98E-03 |  |  |  |  |  |  |
| **Mfge8** | 3.2279 | 1.35E-02 |  |  |  |  |  |  |
| **Il17a** | 14.4229 | 1.35E-02 |  |  |  |  |  |  |
| **Il6** | 7.4833 | 1.35E-02 |  |  |  |  |  |  |
| **C6** | 4.7983 | 1.74E-02 |  |  |  |  |  |  |
| **S100a8** | 12.1584 | 1.96E-02 |  |  |  |  |  |  |
| **Igf1r** | 2.8116 | 2.25E-02 |  |  |  |  |  |  |
| **Rora** | 3.6681 | 2.38E-02 |  |  |  |  |  |  |
| **Ccl22** | 3.8875 | 2.54E-02 |  |  |  |  |  |  |
| **Ptgs2** | 4.8628 | 4.09E-02 |  |  |  |  |  |  |
| **Ccl3** | 4.5947 | 4.27E-02 |  |  |  |  |  |  |

**FIG 3.**
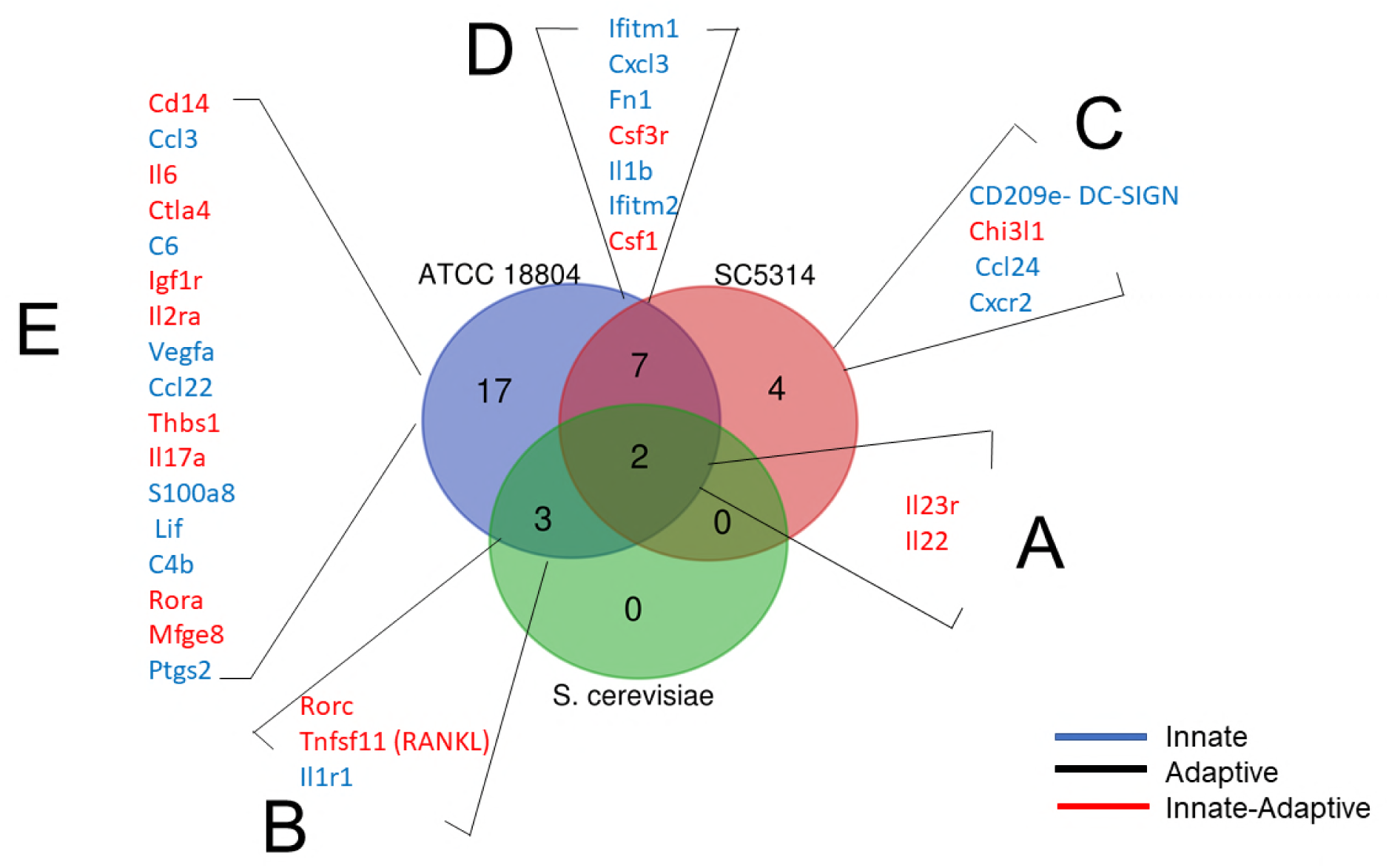
The Venn Diagram shows the differential gene expression of PP CD11c+ phagocytes 24 hrs post-gavage with *C. albicans* strains ATCC18804 and SC5314 and *S. cerevisiae*: **(A)** shared upregulated genes in CD11c^+^ phagocytes when treated with both strains of *C. albicans* and *S. cerevisiae*; **(B)** shared upregulated genes when treated with *C. albicans* ATCC 18804 and *S. cerevisiae*; **(C)** upregulated genes when treated with *C. albicans* SC5314; **(D)** shared upregulated genes when treated with *C. albicans* ATCC 18804 and *C. albicans* SC5314; **(E)** upregulated genes when treated with *C. albicans* SC5314. Genes were also sorted into those that are part of the innate (blue text) or adaptive immune (black text) response and those that play a role in the transition between innate to adaptive immunity (red text).

*C. albicans* strain ATCC18804 and *S. cerevisae* shared the expression of the TNF Receptor Superfamily Member 11b (Tnfs11), the Retinoic Acid-Related Orphan Receptor C (Rorc), and the IL-1 receptor type 1(IL-1r1) **(Fig. 3B and Table 1)**. The Tnfs11 gene was upregulated 3.9-fold for strain ATCC18804 and 5.9-fold upregulation for *S. cerevisiae* Tnfs11 in mice has been suggested to play a role in lymph node organogenesis, as this gene provides the instructions to make osteoprotegerin a protein involved not only in the regulation of osteoclasts, but also in the binding to a protein called receptor activator of NF-κB ligand (RANKL). RANKL has been shown to be an important factor in M-cell differentiation within PPs’ epithelial barrier. M-cells are important in the controlled transcytosis of luminal antigens and microbes. We and others previously reported that *C. albicans*’ translocation into PPs was partially M-cell dependent (16, 17, 27-30). It has been shown that the Rorc deficiency plays a role in susceptibility to *Candida sp.* and *Mycobacterium sp.* Rorc has been shown to be the master gene controlling Th17 differentiation, which is important in immune responses against fungi (31-33). For Rorc, we found a 3-fold upregulation for strain ATCC18804 and 3.7-fold upregulation for *S. cerevisiae.* Research on the IL-1R has shown that signaling plays a critical role in fungal control at the early stages of infection in a mouse model of oropharyngeal candidiasis (OPC). For IL1r1, we observed a 5.1-fold upregulation for strain ATCC18804 and a 4.3-fold upregulation for *S. cerevisiae*. IL1r1 has been found to be important in both the recruitment of circulating neutrophils to the site of infection and the mobilization of newly generated neutrophils from the bone marrow in response to *C*. *albicans* (34).

Treatment with *C. albicans* strain SC5314 upregulated eleven genes in CD11c^+^ phagocytes, some that are part of innate immunity and some that play a role in the transition between the innate and adaptive immune response. Because of this difference, we further divided the genes into two subgroups, those that were not shared with strain ATCC 18804 and those that were shared **(Table 1)**. The four upregulated genes that were not shared with strain ATCC 18804 were Dendritic Cell-Specific ICAM-3-Grabbing Non-Integrin 1 (CD209e, DC-SIGN), Chitinase-like protein 3 (Chil3), C-C Motif Chemokine Ligand 24 (Ccl24), and C-X-C Motif Chemokine Receptor 2 (Cxcr2) **(Fig. 3C)**. The C-type lectin DC-SIGN, which was upregulated 17.5-fold, is an important innate immune receptor expressed on phagocytes such as dendritic cells DCs, and has been demonstrated to recognize N-linked mannans on *C. albicans* (35-37). Chitinase-like protein 3 (Chil3), which was upregulated 31.5-fold, is a pseudo-chitinase that promotes *C. albicans* killing (38). It is expressed by intestinal epithelial cells (IECs) and macrophages in inflamed intestines (39). Chil3 also exhibits dual functions that promote Th2-type inflammation as well as enhancing tissue remodeling and repair (39). Ccl24, also known as myeloid progenitor inhibitory factor 2 (MPIF-2) or eosinophil chemotactic protein 2 (eotaxin-2), which is important in the chemotaxis of eosinophils and basophils, was upregulated 25-fold (40). Cxcr2, which was upregulated 5.5-fold, is a receptor for IL-8, a powerful neutrophil chemotactic factor that also has an important role in the pathogenesis of inflammatory bowel disease (41).

The seven upregulated genes that were shared between SC5314 and ATCC18804 were fibronectin-1(Fn1), Interferon Induced Transmembrane Proteins 1 and 2 (Ifitm1 and 2), C-X-C Motif Chemokine Ligand 3 (Cxcl3), Colony Stimulating Factor 3 Receptor (csf3r), Colony Stimulating Factor 1 (csf1) and IL-1β **(Fig. 3D)**. Fibronectin-1, an extracellular matrix (ECM) protein, was upregulated 41-fold in SC5314 and 16.6-fold in ATCC18804. *C. albicans* is known to have specific fibronectin adhesion molecules that it uses during invasion (42, 43). Ifitm 1 and 2, which were upregulated 5 and 3-fold respectively, have emerged as potent viral inhibitors of a broad range of RNA viruses, although they have been found to be overexpressed in colitis-associated colon cancer and in severely inflamed mucosa of UC and CD patients (44). Cxcl3, which was upregulated 29-fold, plays a role in inflammation and as a chemoattractant for neutrophils that are important in controlling fungal organisms (45). Csf3r, which was upregulated 4.4-fold, has been shown to play a crucial role in the proliferation, differentiation and survival of cells along the neutrophilic lineage (46). Csf1, which was upregulated 5.4-fold, is a cytokine that plays an essential role in the regulation of survival, proliferation and differentiation of hematopoietic precursor cells, especially mononuclear phagocytes such as macrophages and monocytes (47). IL-1β, which was upregulated 3.5-fold, is a proinflammatory cytokine that exerts similar biological activities after interaction with the IL-1R1 activates monocytes, macrophages, and neutrophils (48, 49).

### *C. albicans* strain ATCC 18804 elicits pro-inflammatory genes in PP CD11c^+^ phagocytes

ATCC18804 alone upregulated 17 genes when compared to SC5314 and *S. cerevisiae.* These differentially expressed genes were Vascular Endothelial Growth Factor A (Vegfa), Cytotoxic T lymphocyte antigen–4 (Ctla4), leukemia inhibitory factor (Lif), CD14 Molecule (Cd14), Complement factor 4b (C4b), Thrombospondin 1 (Thbs1), IL-2 receptor alpha (Il2ra), Milk fat globule-EGF factor 8 (Mfge8), IL-17a, IL-6, Complement factor 6 (C6), S100 calcium-binding protein A8(S100a8), insulin-like growth factor 1 receptor(Igf1r), RAR-Related Orphan Receptor A Natural helper (Rora), RAR-Related Orphan Receptor A Natural helper (Ccl22), Prostaglandin-Endoperoxide Synthase 2 (Ptgs2) and C-C Motif Chemokine Ligand 3(Ccl3) **(Table 1; Fig. 3E)**. In examining their functions, the genes that were upregulated have pro-inflammatory functions, as well as protecting the mucosal barrier. Vegfa, which was upregulated 12.7-fold, is a product of CD11c^+^ phagocytes and produces a potent pro-angiogenic mediator that is typically released under hypoxic conditions, but is also a hallmark of inflammation (50). CTLA-4, which was upregulated 6.4-fold, is the major negative regulator of T-cell responses and is also expressed in monocytes and monocyte-derived dendritic cells (DCs) (51). LIF, which was upregulated 6.4-fold, possesses proinflammatory properties in common with tumor necrosis factor-alpha (TNF-α) and IL-1 and −6. LIF also has chemotactic activity through the induction of IL-8 production. LIF has been found in inflammatory bowel diseases, particularly ulcerative colitis (UC). Tissues that have high LIF are histologically characterized by the infiltration of the colonic mucosa with activated neutrophils, macrophages and lymphocytes (52). The CD14 molecule, which was upregulated 6.8-fold, is expressed on monocytes displaying potent antifungal properties such as inhibition of germination and phagocytosis, and can induce a protective Th17 response upon recognition of *C. albicans* (53, 54). As part of the classical complement pathway, complement factor 4b (C4b) was upregulated 4.3-fold. The activation of this factor in the intestinal mucosa is important, as *C. albicans* is able to use C4b-binding protein (C4BP) to inhibit complement activation at the yeast surface that mediates adhesion of *C. albicans* to host cells (55, 56). Extracellular matrix protein thrombospondin-1 (Thbs1), which was upregulated 11-fold, is rapidly and transiently secreted at high concentrations by macrophages, endothelial cells, and fibroblasts, and is released at sites of tissue injury and inflammation. The effects of TSP-1 include, but are not limited to, neutrophil and monocyte chemotaxis phagocytosis of neutrophils, regulation of T cell function via receptor ligation, activation of latent TGF-β1, and the inhibition of angiogenesis (57). It has also been shown that Thbs1 can also enhances the pathogenesis of disseminated candidiasis by creating an imbalance in the host immune response that ultimately leads to reduced phagocytic function, impaired fungal clearance, and increased mortality (58). The milk fat globule-EGF factor 8 protein (Mfge8), which was upregulated 3.2-fold, encodes a protein product, lactadherin, a membrane glycoprotein that promotes phagocytosis of apoptotic cells. Mfge8 has been shown to play an important role in maintaining the integrity of the intestinal mucosa and to accelerate healing of the mucosa in septic mice by the phagocytic removal of damaged and apoptotic cells from tissues, the induction of VEGF-mediated neovascularization, the maintenance of intestinal epithelial homeostasis, and the promotion of mucosal healing (59, 60). The well-studied and important interleukin in fungal infections, Il-17a, was upregulated 14.3-fold. Although IL-17 was initially thought to be produced almost exclusively by T cells, it has lately become clear that this cytokine is also made by a variety of innate cell types such as γδ-T cells, NKT cells, lymphoid tissue inducer (LTi) cells, macrophages, and possibly DCs and neutrophils (61). IL-17’s specific role is in the protection against the fungus *C. albicans*. The IL-17 pathway regulates antifungal immunity through upregulation of pro-inflammatory cytokines including IL-6 and neutrophil-recruiting chemokines such as CXCL1 and CXCL5 (62). IL-6, which was upregulated 7.5-fold, is a pro-inflammatory cytokine found in the early stages of *C. albicans* infection (63). Monocytes and macrophages are the main sources of Il-6, and it has been found that IL-6 deficient mice are particularly susceptible to systemic candidiasis (64, 65). The complement C6 (C6) gene, which was upregulated 4.8-fold, produces the C6 protein important in forming the membrane attack complex (MAC), which plays an important part in the defense against ATCC 18804 by augmenting phagocytosis (56, 66). S100 calcium-binding protein A8 (S100a8), which functions as an alarmin, was upregulated 12.1-fold. S100a8 is pro-inflammatory that is enhanced by the presence of IL-17. S100 proteins are important in PMN neutrophil responses (61, 67-70). The insulin-like growth factor 1 receptor (Igf1r), which was upregulated 2.8-fold, is important in augmenting polymorphonuclear neutrophilic leukocytes (PMNLs) during Candida infection (71). Produced by natural helper (NH) cells, RAR-Related Orphan Receptor A (Rora), which was upregulated 3.6-fold, is an innate lymphoid cell (ILC) that produces T helper-2 (Th2)-cell-type cytokines in the lung-and gut-associated lymphoid tissues. (72). Chemokine C-C Motif Chemokine Ligand 22 (Ccl22), which was upregulated 3.8-fold, plays an important in the chemotactic for monocytes, dendritic cells and natural killer cells (73). Prostaglandin-endoperoxide synthase-2 (Ptgs2), also known as cyclooxygenase, was upregulated 4.8-fold. Ptgs2 is the key enzyme in prostaglandin biosynthesis, and acts both as a dioxygenase and as a peroxidase. It has been found that in *C. albicans*-infected resident peritoneal macrophages, activation of group IV RprcA cytosolic phospholipase A2 (cPLA2α) by calcium- and mitogen-activated protein kinases triggers the rapid production of prostaglandins I2 and E2 through cyclooxygenase (COX)-1 (74). The Chemokine (CC-motif) ligand 3 (CCL3), which was upregulated 4.6-fold, has been shown to play important roles in the acute inflammatory response to *C. albicans* corneal infection (75) **(Fig. 3E;Table 1)**.

To our surprise, the differential gene expression of the Dectin-1 receptor in PP CD11c^+^ phagocytes did not meet our stringent cut-off criteria. To test the fold change in gene expression for Dectin-1, we performed real-time PCR. The results revealed a 2.5-fold change for Dectin-1 in PP CD11c^+^ phagocytes that had been gavaged with strain SC5314, compared to the 1-fold gene expression change for strain ATCC 18804 **(Fig S2).** One reason we could not detect Dectin-1 by nanostring may be explained by the work of Vautier and colleagues, who demonstrated that Dectin-1 was not required for controlling *C. albicans* colonization of the gastrointestinal tract (76).

### Confirmation of upregulated gene expression profiles of CD11c^+^ phagocytes by CBA and real-time PCR

To test if we could recapitulate some of the nanostring results at the protein cytokine level, we used the cytometric bead array assay that allows for the measurement of multiple soluble proteins at one time by flow cytometry (77, 78). The advantage of CBA over ELISA is that it requires a small amount of sample, which is ideal for working with PP single-cell suspensions. To do this, we harvested PPs from mice that had been gavaged with *C. albicans* strains SC5314, ATCC18804, and *S. cerevisiae*. Total PP cells were lysed and the IL-1β, IL-6, IL-17f, IL-17a and IL-22 protein cytokines were measured using a cytometric bead array assay. The results reveal that PPs that had sampled *C. albicans* strain ATCC 18804 had elevated protein cytokine levels that were 2-3-fold higher than PPs that sampled SC5314, *S. cerevisiae* or PBS, suggesting that the ATCC 18804 is recognized as a potential danger by PPs **(Fig. 4)**.

**FIG 4.**
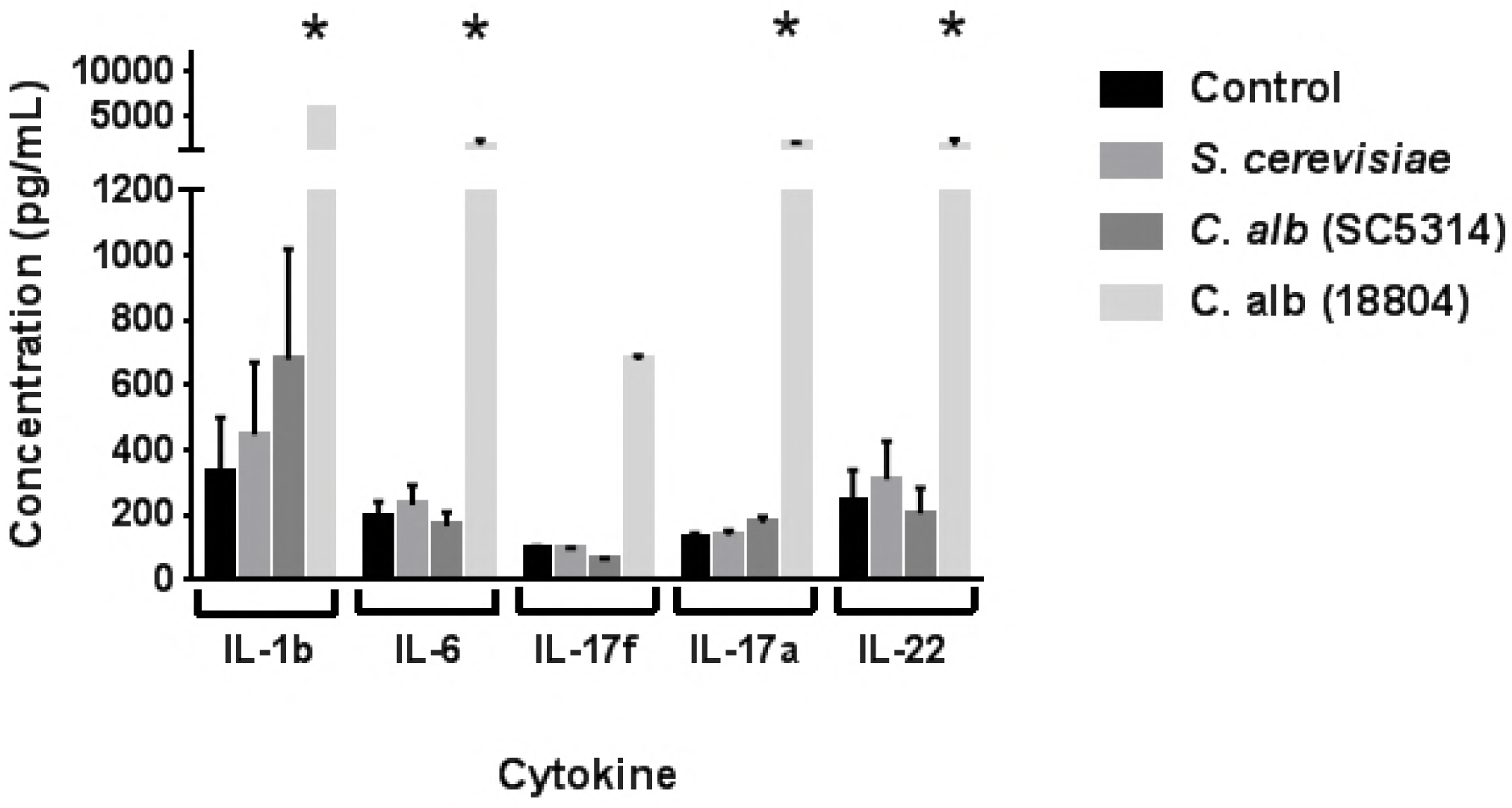
Cytometric bead array assay recapitulates nanostring results at the protein cytokine level. PPs from mice that had been gavaged with *C. albicans* strains SC5314, ATCC18804 and *S. cerevisiae* were harvested and lysed. Protein cytokines IL-1b, IL-6, IL-17f, IL-17a and IL-22 were measured using cytometric bead array. PPs that had sampled *C. albicans* strain ATCC 18804 had elevated protein cytokine levels that were 2 to 3-fold higher than PPs that sampled *C. albicans* strain SC5314, *S. cerevisiae* or PBS. Statistical significance p< 0.05.

In addition to testing upregulation of five cytokines by CBA in PP, we confirmed additional upregulated factors like Cd14, Fn1, S100a8 and Thbs1 using reverse transcription RT-qPCR. Similar to the Nanostring data, the RT-qPCR results reveal that in *C. albicans* 18804-treated CD11c^+^ phagocytes, Cd14 was upregulated 3-fold, Fn1 was upregulated 10-fold, S100a8 was upregulated 4-fold and Thbs1 was upregulated 10-fold. The data were normalized to housekeeping gene Tubb5 (**Fig. 5).** We also tested another unchanged housekeeping gene, Hprt, that showed no change in expression in RT-qPCR assay. The results obtained by CBA and RT-qPCR methods demonstrated that our nanostring data were reproducible.

**FIG 5:**
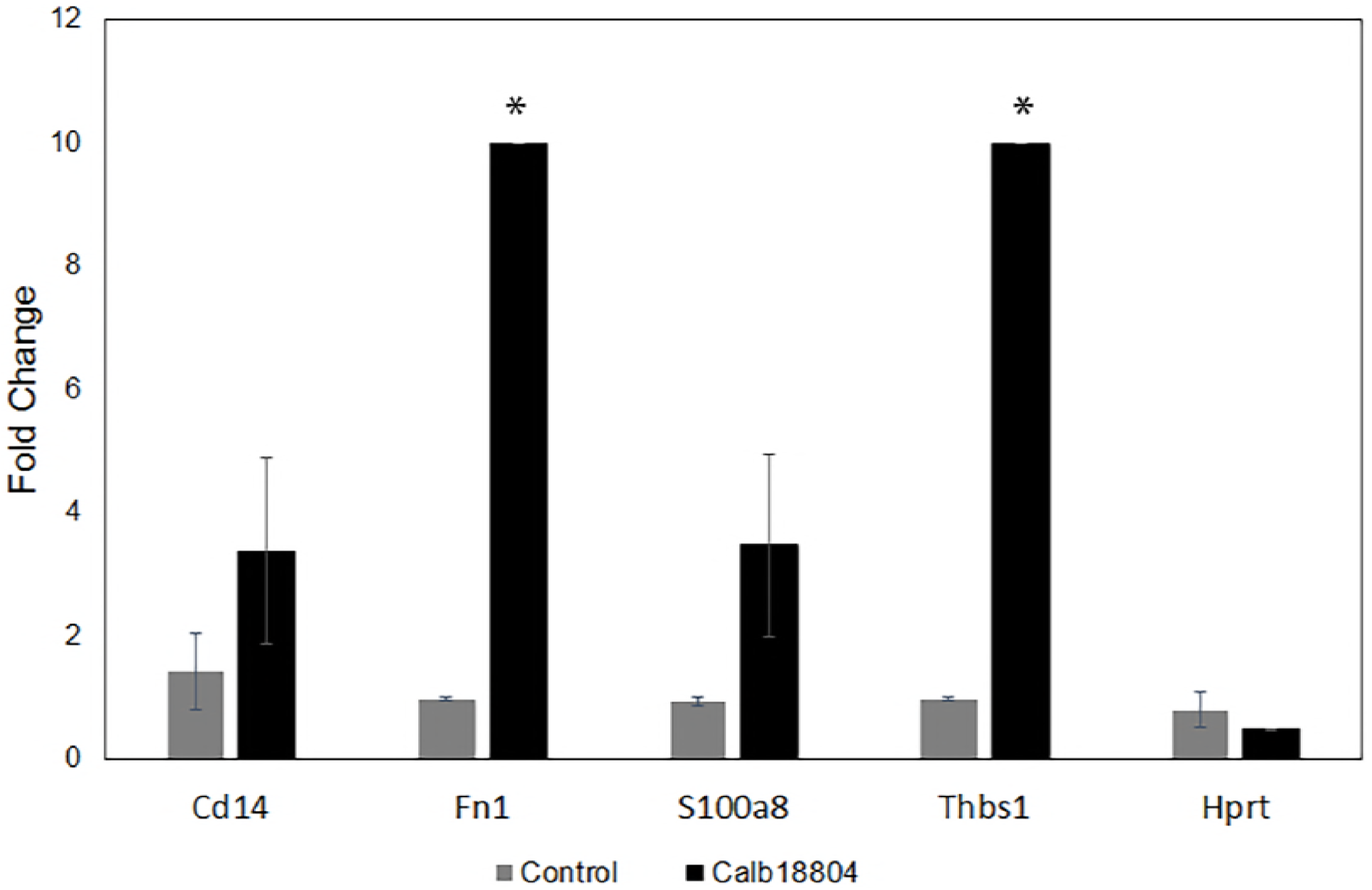
A subset of significantly upregulated genes in sorted PP CD11c^+^ DC phagocytes that have sampled C. albicans 18804. Genes that were in the nanostring panel were independently tested by quantitative real-time PCR using SYBR Green assay. Histogram shows the fold enrichment of 4 upregulated genes (Cd14, Fn1, S100a8, Thbs1) and one unchanged gene, Hprt, were normalized relative to Tubb5 control. The experiment was performed as replicates by pooling cells from 3 mice each in control and treatment groups. Statistical significance p< 0.05.

## Discussion

It is estimated that fungi represent less than 0.1% of the total microbes in our gut microbiota; however, due to an incomplete fungal genomic database, this is likely to be an underestimate (79-81). Recent efforts have started to elucidate the composition of the fungal mycobiome, and based on these studies, we know that the GI tract has the largest composition of the *Candida* genus, which is comprised of about 160 species (4, 79, 82, 83). Although *Candida* is part of the microbiome, recent studies have also shown that altered bacterial diversity and an increased fungal load of *Candida* species is associated with IBD, particularly Crohn’s disease (81, 84). There is still a lot of fascinating work to be done in this area to characterize the mycobiome and understand how dysbiosis leads to human gastrointestinal disease. However, little is known about the initial contact between *Candida* and the gut-associated lymphoid tissues during sampling and immune surveillance. Understanding which cell types respond to the presence of *C. albicans in vivo* is important because this will allow for the identification of potential therapeutic targets and determine which cell types might be predictors of *Candida*-associated GI diseases.

To address how GALT such as Peyer’s patches detect *C. albicans*, and whether important immune responses were elicited, we chose to determine gene expression profiles within the mouse PPs during the first 24 hours of sampling. In our model, we gavaged mice with two strains of *C. albicans*, ATCC18804 and SC5314, because it has been shown that the innate immune receptor Dectin-1 receptor is essential for controlling infection in *C. albicans* strain SC5314, but is not required for *C. albicans* strain ATCC 18804. This difference in immune activation made these strains good candidates for study to understand whether PPs were able to discriminate between them and the *S. cerevisiae* control.

In a previous study, we were able to isolate PP-specific CD11c^+^ phagocytes, CD3^+^ T-cells, and B220^+^ positive B cells by cell sorting, as these three major cell types are critical in PP sampling and immune surveillance **(Fig. 6)**. We were also inspired by a previous study by Xu et al., where gene expression profiling of *C. albicans* was done during infection using the nanostring and counter platform (82). We also chose nanostring technology because unlike RNAseq, the targeted transcriptomics do not require library preparation and complex data processing. Gene expression profiling of PPs that sampled *C. albicans* and *S. cerevisiae* showed that these tissues are able to discriminate between strains and other fungal organisms. The data revealed that CD11c-positive phagocytes are the specific cell type that shows differential gene expression within 24 hours of sampling. This matches with our confocal microscopy data, as we have detected sampling at this time point. This does not mean that that B and T lymphocytes do not have differential gene expression; rather, at 24 hr time point, phagocytes are the cell types that sample and coordinate the signals that activate innate and adaptive immune responses.

**FIG 6.**
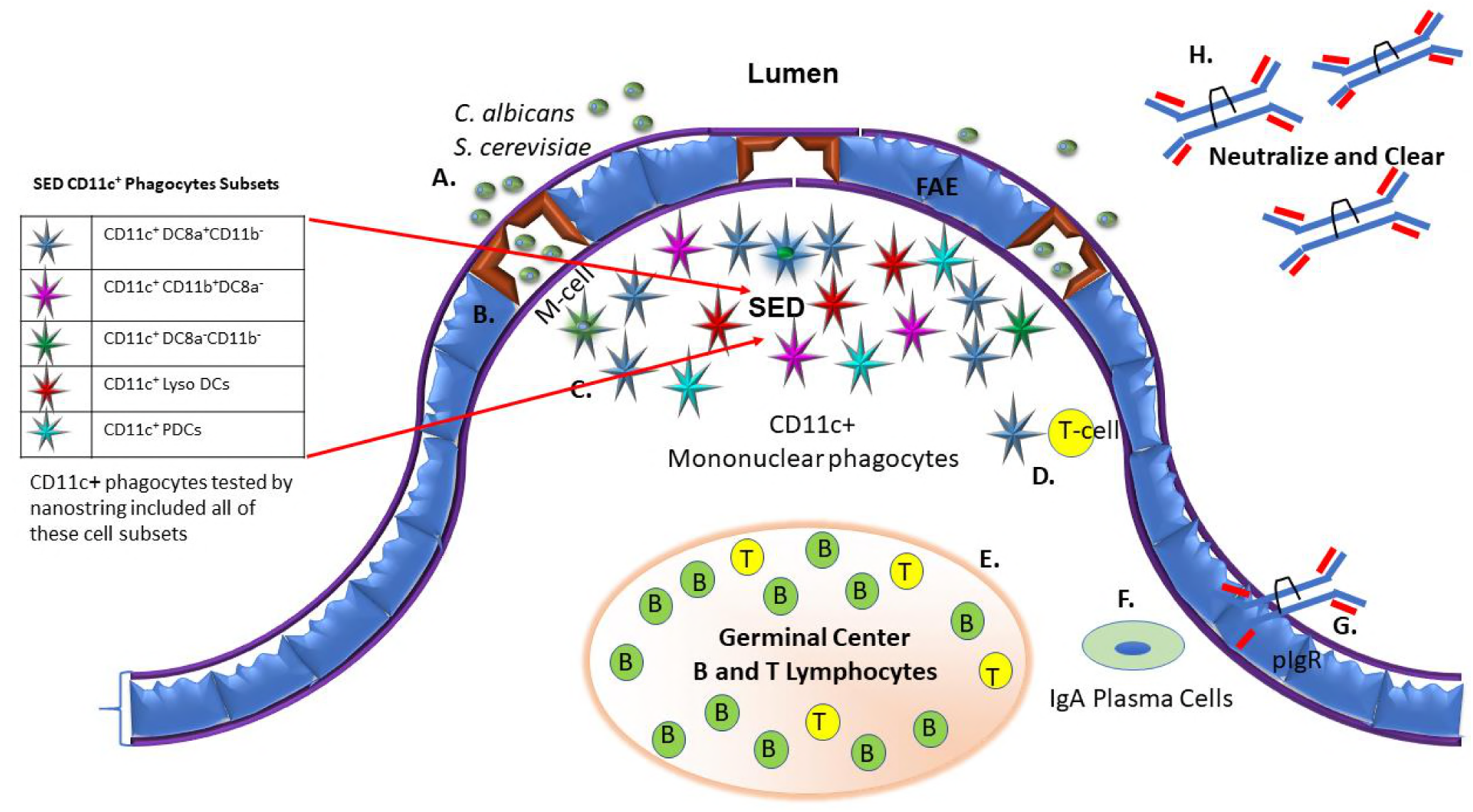
Sampling and trans-epithelial transport of antigen and microbes by Peyer’s patches. (A) Antigen/microbes adhere to the apical surfaces of M-cells. (B) Antigens/microbes are endocytosed by M cells and delivered into the M cell pocket. (C) Antigens/microbes can also interact with the five known CD11c^+^ phagocyte subsets that reside in the sub-epithelial dome (SED). This is the site where differential gene expression occurs within the first 24 hrs post-gavage with *C. albicans* strains ATCC18804, SC5314 and *S. cerevisiae* (red arrow). (D) DCs can then present to resident follicular T-cells which reside in the intrafollicular regions (IFR) and secrete cytokines that promote B cells in germinal center (GC) to undergo class-switch recombination (CSR) and somatic hypermutation (SHM). (E) B cells migrate into the mucosa through high endothelial venules (HEVs) and into the lamina propia. (F) Activated B plasma cells in produce polymeric IgA that are exported as SIgA via the pIgR receptor. (G) Once in the lumen, SIgA promotes the clearance of antigens and microbes.

We also found that both strains of *C. albicans*, as well as *S. cerevisiae*, stimulate two important genes in CD11c^+^ phagocytes, IL-22 and IL-23. These cytokines are known to play important roles in maintaining barrier integrity and antimicrobial activity, as well as stimulating tissue-damage protection, tissue repair and remodeling, and antifungal immunity. This suggests that PPs are able to detect fungi as foreign and are prepared to defend the mucosal barrier. In contrast to studies by Bonifazi et al., we observed that these protective immune responses do not require the hyphal form of *C. albicans* (18). We found that some genes were shared between *C. albicans* strain ATCC18804 and *S. cerevisae, C. albicans* strain SC5314, and *S. cerevisae and C. albicans* strains ATCC18804 and SC5314. All of these genes were important in the innate immune response as well as the transition from innate immunity to adaptive. *S. cerevisae* did not stimulate its own standalone set of genes, but it did stimulate the expression of the Tnfs11, also known as RANKL, which has been shown to be an important factor in M-cell differentiation within the PPs epithelial barrier. However, it is unclear whether the purpose of increasing RANKL gene expression is to stimulate more sampling. The Retinoic Acid-Related Orphan Receptor C (Rorc) and IL-1r1 were also upregulated, and these play an important role in fungal control at the early stages of infection.

We also found that *C. albicans* strain SC5314 stimulated important lectins, DC-SIGN that recognizes N-linked mannans on *C. albicans* and Chil3, which not only promotes *C. albicans* killing, but also promotes Th2-type inflammation, as well as enhancing tissue remodeling and repair. *C. albicans* strain SC5314 also stimulates other genes that are important in chemotaxis and neutrophil recruitment such as Ccl24, also known (eotaxin-2), and Cxcr2. Cxcr2 a powerful neutrophil chemotactic factor that also has an important role in the pathogenesis of inflammatory bowel disease. To our surprise, we did not detect Dectin-1 when using nanostring technology, but when we performed real-time PCR, we found a modest 2.5-fold change for Dectin-1 in PP CD11c^+^ phagocytes that had been gavaged with strain SC5314. A possible reason why we could not detect Dectin-1 by nanostring technology may be explained by Vautier and colleagues, whose work demonstrated that Dectin-1 was not required for controlling *C. albicans* colonization of the gastrointestinal tract and may be redundant (76). Our data suggests that other chemokines, cytokines and innate immune receptors that are stimulated by CD11c^+^ phagocytes may be the reason why Dectin-1 in the gut is dispensable. SC5314 also shared genes with ATCC18804 that were pro-inflammatory, such as Fn1, Ifitm1 and 2, Cxcl3, csf3r, csf1 and Il-1β. Interestingly, Fn1, an extracellular matrix (ECM) protein, was upregulated 41.1-fold. *C. albicans* is known to have specific fibronectin adhesion molecules that it uses to exploit fibronectin during invasion and possibly in the gut. However, another function of fibronectin is that it can influence the maturation of dendritic cell progenitors (42, 43, 85).

*C. albicans* strain ATCC18804 not only upregulates genes in CD11c^+^ phagocytes involved in pro-inflammation, such as Vegfa, Ctla4, Lif, CD14, C4b, Thbs1, IL-2rα, C6, S100a8, Igf1r, Rora, Ccl22, Ptgs2, and Ccl3, but it also stimulates genes that are important in the specific control of fungi, such as Il-17a and IL-6. Interestingly, ATCC18804 also upregulated the MFG-E8 gene, which has been shown to play an important role in maintaining the integrity of the intestinal mucosa and to accelerate healing of the mucosa in septic mice by the phagocytic removal of damaged and apoptotic cells from tissues, leading to the maintenance of intestinal epithelial homeostasis. This data suggests that although mucosal tissues sample potentially pathogenic organisms, this tissue is also quite dynamic, as immune cells are also sending signals to repair and to restore homeostasis. We were also able to show that these changes in gene expression translate at the protein level, and that strain ATCC 18804 was able to stimulate strong protein cytokine responses in PPs.

In summary, by interrogating the PPs transcriptome, we discovered that PPs can identify and respond to fungi such as *C. albicans* and *S. cerevisiae*. We demonstrated that within the first 24 hours of sampling, CD11c^+^ phagocytes were not only important in sampling, but they were the cell type that exhibited differential gene expression. Our results also revealed that PPs can distinguish between different strains of *C. albicans*. Interestingly, the differentially expressed genes play important roles inflammation, chemotaxis and fungal-specific recognition, as well as maintaining homeostasis and protection of the mucosal barrier. Our future work will use RNAseq to address temporal and comprehensive changes in gene expression of PP, CD11c^+^ phagocytes, B220^+^ B-cells, and CD3^+^ T-cells in order to map shifts in gene expression from innate to adaptive immunity. We found that using PPs cell subsets is informative and can be applied universally to study how the intestinal mucosa recognizes a variety of organisms.

## Materials and Methods

### Animals

Swiss Webster female mice (8–12 weeks old) were obtained from Taconic Farms (Hudson, NY). Animals were housed under conventional, specific pathogen-free conditions and were treated in compliance with the Wadsworth Center’s Institutional Animal Care and Use Committee (IACUC) guidelines.

### Animal Use Ethics Statement

Studies that involved mice were reviewed and approved by the Wadsworth Center’s IACUC under protocol #18-450. The Wadsworth Center complies with the Public Health Service Policy on Humane Care and Use of Laboratory Animals and was issued assurance number A3183-01. The Wadsworth Center is fully accredited by the Association for Assessment and Accreditation of Laboratory Animal Care (AAALAC).

### Maintaining RNA integrity during PP excision for Nanostring

Briefly, mice were gavaged with 1×10^8^ *C. albicans* strains ATCC 18804, SC5314 or *Saccharomyces cerevisiae* strain in a total volume of 100 µl using a 22 G ×1.5 in. blunt-end feeding needle. Mice were sacrificed 24 hrs post-gavage and PPs were excised and placed into 2mls of cold HBSS containing 50 µl of SUPERase·In RNase inhibitor (Thermo Fisher, Waltham, MA). PPs were transferred into 2mls of fresh HBSS that contained 30 µl of SUPERase·In RNase inhibitor.

### Cell Sorting and RNA isolation from PPs cells

PP single-cell suspensions were made by grinding PPs through a 70 µm mesh filter (BD Bioscience, San Jose, CA). Cells were transferred to a 96 well U-bottom plate and Fc receptors on cells were blocked with Rat anti-mouse Fc-Block CD16/CD32-Clone 2.4G2 (BD Pharmigen, San Jose, CA) on ice for 15 minutes. PP cells were then stained with a cocktail of B-cell, 20 µg /ml Rat anti-mouse CD45R/B220 PerCp Clone RA3-6B2 (BD Pharmigen, San Jose, CA), 50 µg /ml T-cell, Rat anti-mouse CD3 FITCClone 17A2 (BD Pharmigen, San Jose, CA), and 20 µg /ml CD11c^+^ phagocytes Armenian Hamster anti-mouse CD11cClone N418 (eBioscience, San Diego, CA) on ice for 30 min as previously described (12). B and T-lymphocytes and CD11c^+^ phagocytes were sorted on a FACSAria IIu Flow Cytometry Sorter (BD Biosciences, San Jose, CA) and into 0.22 µm Spin X centrifuge tube filters (Corning, Tewksbury, MA) that were pre-moistened with 100 µl of flow buffer that also contained SUPERase·In RNase inhibitor. For PP B and T-cells, 300,000 cells were sorted for each cell type in individual Spin X tubes. Due to their complex structure and sparse numbers of DCs, we sorted 50,000-80,000 cells. After sorting, the Spin-X column was centrifuged at 2,000 rpm for 2 min to ensure that all cells were collected into the filter. The cells were then lysed on the filters by increasing the centrifugation speed to 13,400 rpm for 1 min. RNA was isolated on the Spin X column using RNA Easy Kit (Qiagen, Netherlands) and the DNAse RNase-Free Set (Qiagen, Netherlands) according to manufacturer’s instructions and as described previously (12). To ensure that the RNA quantity and quality were sufficient, isolated RNA was measured with an Agilent RNA 6000 Pico kit following the protocol in the manufacturer’s instructions.

### Measuring RNA counts by Nanostring

Since RNA yields for PP cells range between 1-20 ng, the nCounter Low RNA Input Amplification kit was used to generate cDNA. A multiplexed target enrichment (MTE) primer pool was used to amplify target genes in the code set prior to hybridization with the nCounter reporter and capture probe set. The desired amplification of targets in all cell types was checked by reverse transcription PCR and by RT-qPCR before proceeding with nCounter Low RNA Input amplification. After amplification, RNA counts were assessed with the nCounter® mouse Pan Cancer Immune profiling kit that profiles over 770 murine immune system-related genes. A detailed protocol for PPs using this method was previously described (12).

### Staining of PPs for Confocal Microscopy

PP from mice that were gavaged with FITC-labeled *albicans* strains 18804 and SC5314 and *Saccharomyces cerevisiae* were harvested 24 hrs post-gavage, snap-frozen in liquid nitrogen, and embedded in TissueTek Optimal Cutting Temperature Compound (OCT; Sakura Finetek, Torrance, CA). 20 µm of serial cryosections were prepared with a Themo-Fisher NX-70 cryostat (Thermo Fisher, Waltham, MA). Cryosections were air-dried at room temperature for 24 h before immunostaining. PPs were then fixed for 2 min in acetone, washed in PBS-Tween 20 (PBS-T); 0.05%, encircled with an ImmEdge hydrophobic pen (Vector Labs, Burlingame, CA), and blocked with 2% fetal calf serum in 1X PBS for 30 min at 37°C. Slides were rinsed for 10 sec in PBS containing 0.5% Tween 20 (PBS-T), and Fc receptors were then blocked for 15 min at 37°C with supernatants from the ATCC 2.4.G2 hybridoma cells, which secrete an antibody against Fc gamma receptors FcRII (CD32). Sections were stained with 10 µg of CD11c-PE for 30 minutes and washed twice with 1X PBS-T. Sections were washed once with PBS-T for 3 min, dried and mounted with ProLong Gold Antifade reagent (Invitrogen, Waltham, MA).

### Microscopy and image processing

Images were captured using a Leica SP5 confocal laser scanning microscope (Leica, Wetzlar, Germany). Panels containing confocal images were generated using Adobe Photoshop version 13.0 × 32 (San Jose, CA). Images were marked using the drawing tools to highlight the results and to provide orientation of PP tissues.

### Real-time PCR

A real-time PCR assay was performed on ABI 7500 FAST Sequence Detection System for Cd14, Fn1, S1008a8, Thbs1, and two housekeeping genes, Hprt and Tubb5. RNA was isolated from CD11c^+^ phagocytes from mice that were gavaged with control and *C. albicans* strain ATCC 18804. The RNA was DNase treated with TURBO DNA-free kit (Invitrogen) prior to cDNA synthesis. cDNA synthesis was performed using Protoscript II First Strand cDNA Synthesis kit (NEB) using random primers. cDNA was then used with Luna Universal qPCR Master Mix and primers listed in **Table 2**. The PCR conditions were 95^°^C for 60 sec, 95^°^C for 15 sec, and 60^°^C for 30 sec for 40 cycles. The fold change in expression was calculated using ddCT method. The Tubb5 was used as normalization control.

**Table 2.**
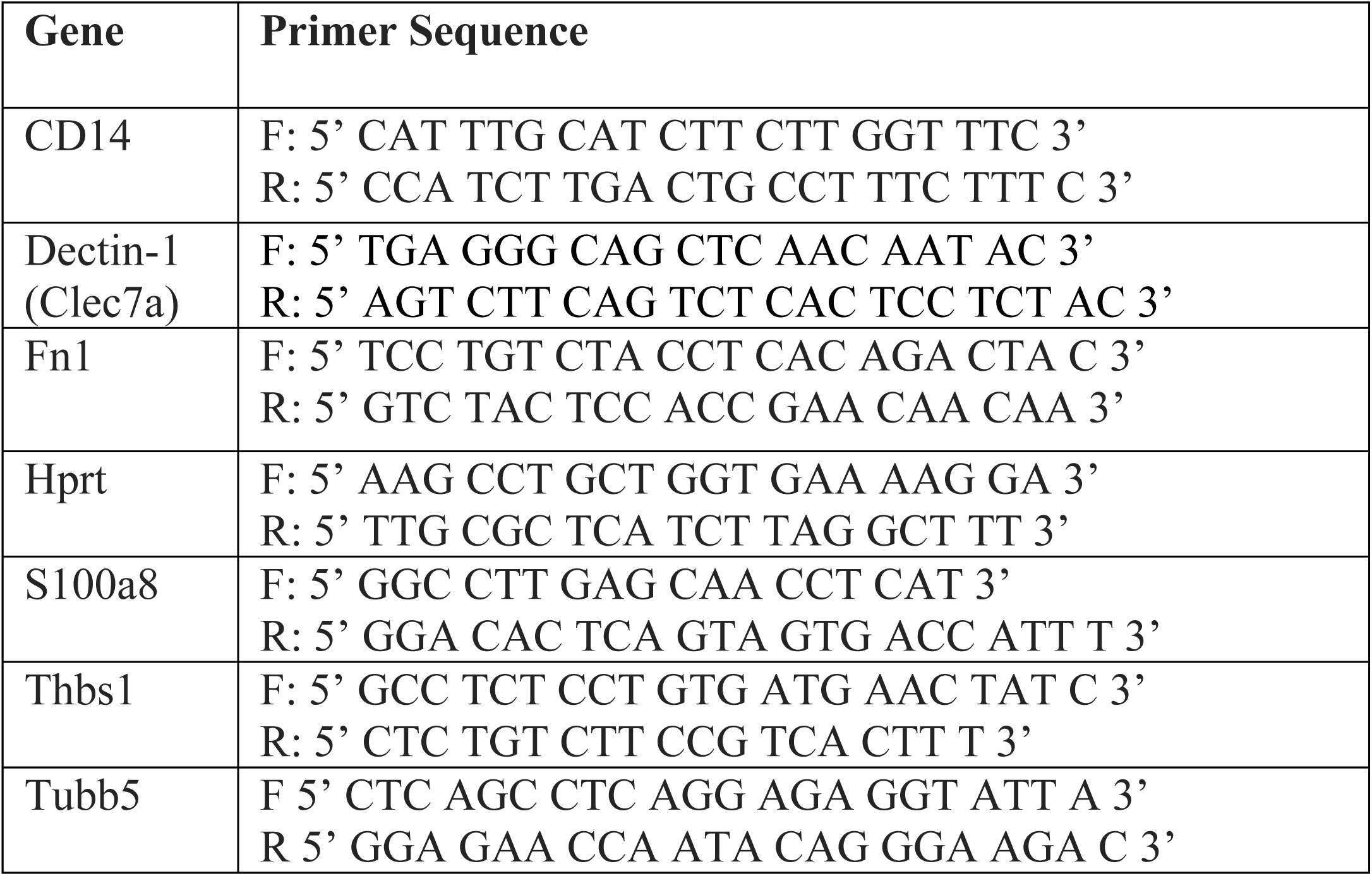
Primers used for RT-PCR

### Cytokine measurements in PP CD11c^+^ phagocytes

CD11c^+^ phagocytes from mice gavaged with *C. albicans* strains 18804 and SC5314 and *Saccharomyces cerevisiae* were sorted into 1X PBS pH 7.4. Cells were spun down at 13,400 rpm to remove the supernatant. NP40 cell lysis buffer containing the addition of 0.001M PMSF and 10X protease inhibitor (Thermo Fisher, Waltham, MA) was added to the sorted cells, and the tube was placed on dry ice for two min. Cells were allowed to thaw for two min followed by an additional 10 min incubation on dry ice. Cells were then allowed to thaw and then placed back on ice for an additional 15 mins. The protein extracts were used at a 1:2 dilution to measure the cytokines IL-22, IL-17A, IL-17F, IL-1β and IL-6 using the LEGENDplex multi-analyte flow assay kit (Biolegend, San Diego, CA) according to manufacturer’s instructions. The cytokine detection beads were measured on a FACS Calibur (BD Biosciences, San Jose, CA).

### Statistical analysis

NanoString analysis was performed in duplicate per experimental condition with the nCounter Analysis System using PanCancer Immune profiling kit. The results were analyzed using raw counts with DEseq2 (86, 87). A transcript was considered differentially expressed when it was up or downregulated at least 2-fold and had a *P*-adj value of <=0.05.

### Data availability

The that authors make the data presented in in this manuscript fully available and without restriction upon request.

### Supplemental material

Supplemental material for this article may be found at URL.

## Acknowledgements

We thank the Wadsworth Center Applied Genomic Technologies Core and the Wadsworth Center Biochemistry and Immunology Core. We thank Dr. Kelly Miller of Nanostring Technologies for assistance with technical support and data analysis. We thank Yannick David of Union College for his help in generating the PP model. We thank Dr. Sudha Chaturvedi at the Wadsworth Center Mycology laboratory for providing *C. albicans* strain ATCC 18804. We thank Dr. Aaron Mitchell at Carnegie Mellon University for providing *C. albicans* strain SC5314. We thank Dr. Kimberly McClive-Reed of Health Research, Inc. for critical reading of this manuscript. M.D.J. was supported by the University at Albany and Wadsworth Center start-up funds. The funders had no role in study design, data collection and interpretation, or the decision to submit the work for publication.

---

**FIG S1:**
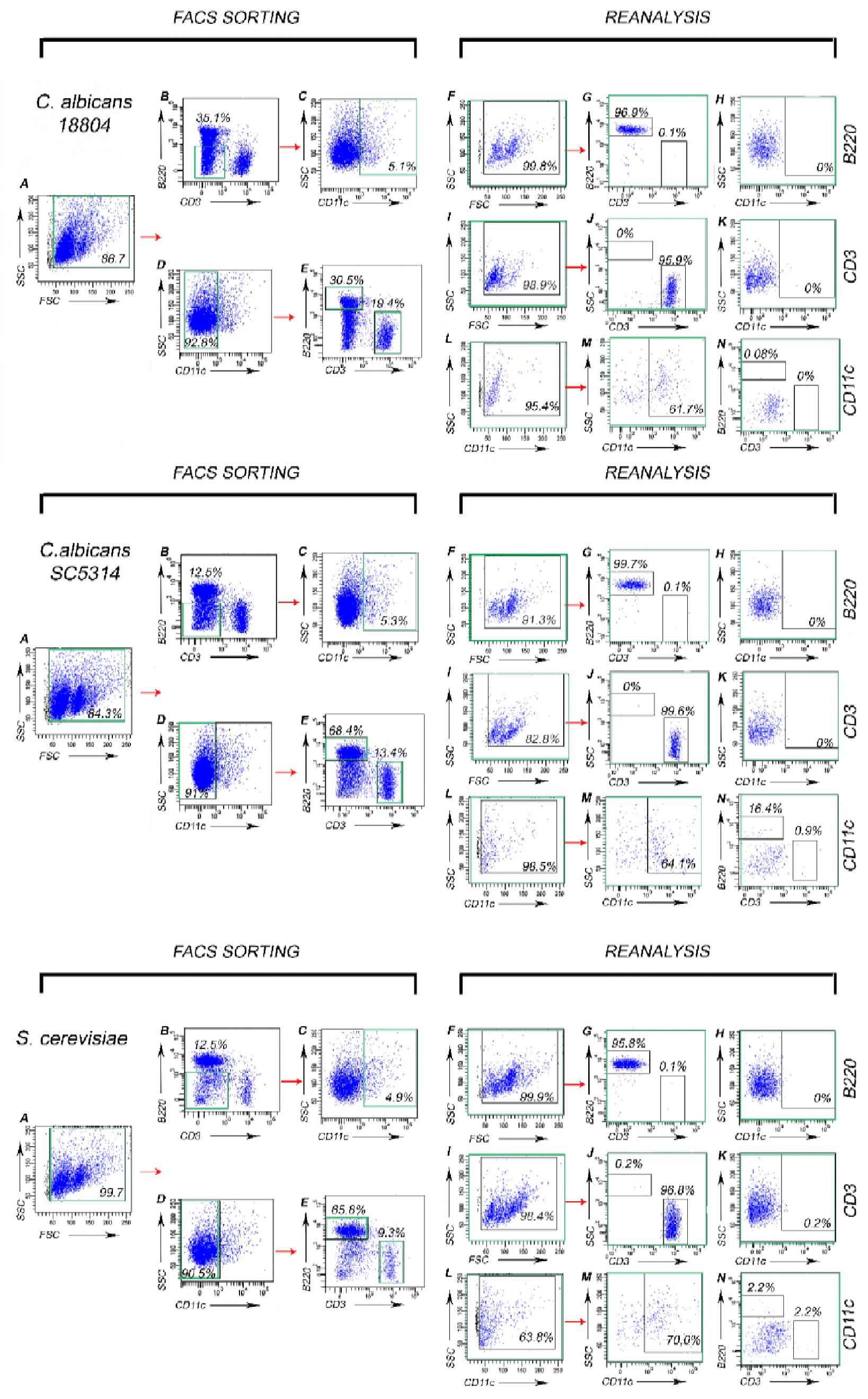
Cell Sorting and Reanalysis of Isolated PP B-cells, T-cells and CD11c^+^ phagocytes. Mice were gavaged with *C. albicans* strains ATCC18804, SC5314 and *S. cerevisiae*, and PPs were harvested 24 hrs post gavage. Single-cell suspension of PPs were stained for the cell surface markers: B220 for B-cells, CD3 for T-cells and CD11c phagocytes. **FACS Sorting Panels:** (A) Forward (FSC) and side scatter (SSC) plots represent the entire single cell suspension. (B) We gated on B220^-^/CD3^-^ population for each treatment group. (C) We gated on only CD11c^+^ phagocytes for each treatment group (D) We gated on cells that were CD11c^-^. (E) We gated on B220^+^ CD3^-^ and B220^-^CD3^+^. **Reanalysis Panels:** (F-H) show the purity for each cell type for B220^+^. (I-K) show the purity for CD3^+^ T-cells. Both B an T cells did not have contamination with CD11c+ phagocytes. (L-N) CD11c^+^ phagocytes were sorted such that these did not have contamination with B or T-cells.

**FIG S2.**
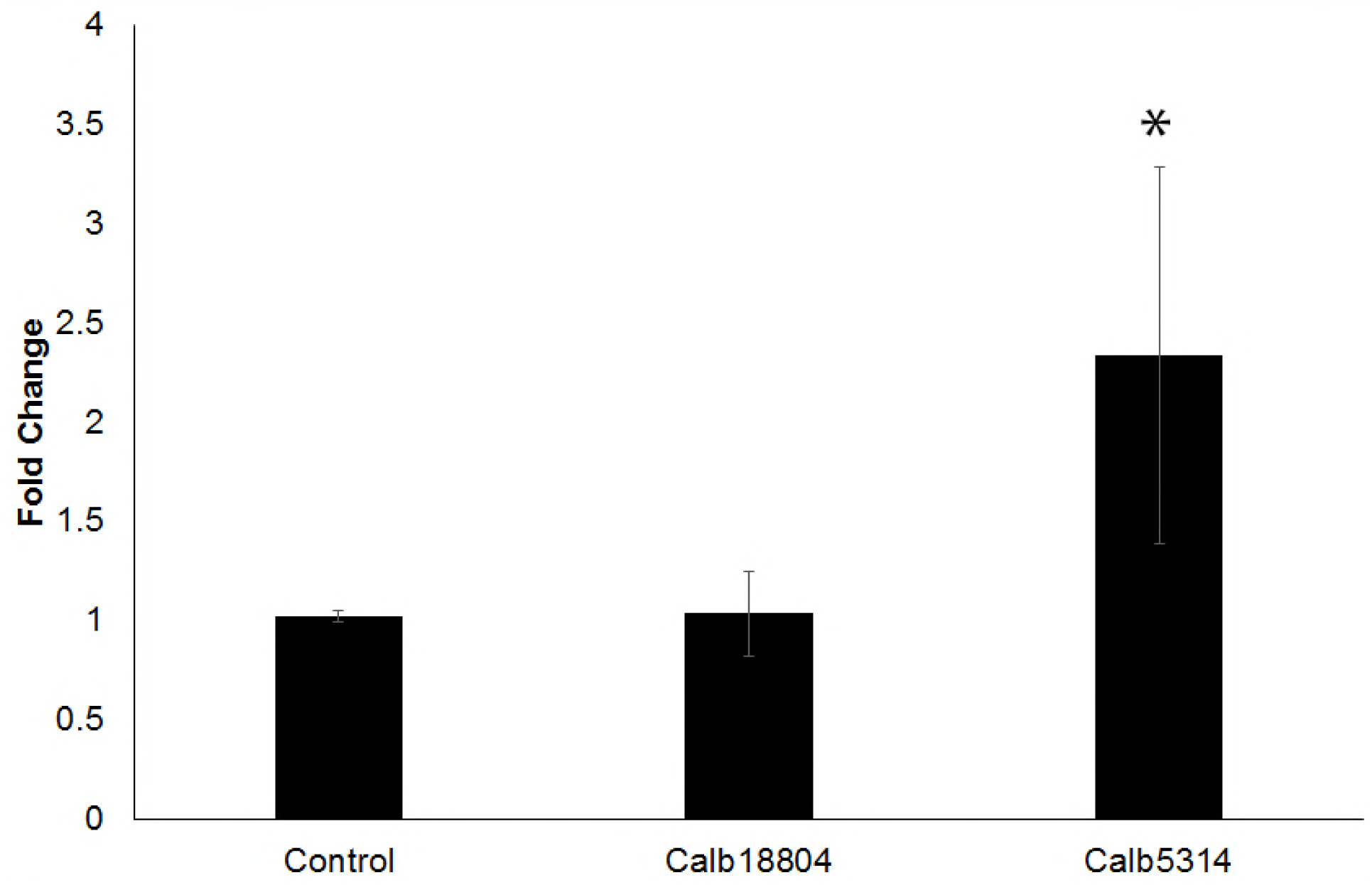
Gene expression of the Dectin-1 receptor in PP CD11c+ phagocytes tested by real-time PCR. A 2.5-fold change was observed for Dectin-1 expression in PP CD11c^+^ phagocytes that had been gavaged with strain SC5314, compared to a 1-fold gene expression change for strain ATCC 18804 and the PBS control. Statistical significance p< 0.05.

